# Distinct error rates for reference and non-reference genotypes estimated by pedigree analysis

**DOI:** 10.1101/2020.02.06.937649

**Authors:** Richard J. Wang, Predrag Radivojac, Matthew W. Hahn

## Abstract

Errors in genotype calling can have perverse effects on genetic analyses, confounding association studies and obscuring rare variants. Analyses now routinely incorporate error rates to control for spurious findings. However, reliable estimates of the error rate can be difficult to obtain because of their variance between studies. Most studies also report only a single estimate of the error rate even though genotypes can be miscalled in more than one way. Here, we report a method for estimating the rates at which different types of genotyping errors occur at biallelic loci using pedigree information. Our method identifies potential genotyping errors by exploiting instances where the haplotypic phase has not been faithfully transmitted. The expected frequency of inconsistent phase depends on the combination of genotypes in a pedigree and the probability of miscalling each genotype. We develop a model that uses the differences in these frequencies to estimate rates for different types of genotype error. Simulations show that our method accurately estimates these error rates in a variety of scenarios. We apply this method to a dataset from the whole-genome sequencing of owl monkeys (*Aotus nancymaae*) in three-generation pedigrees. We find significant differences between estimates for different types of genotyping error, with the most common being homozygous reference sites miscalled as heterozygous and vice versa. The approach we describe is applicable to any set of genotypes where haplotypic phase can reliably be called, and should prove useful in helping to control for false discoveries.

## Introduction

When dealing with large sets of genotype data, such as those obtained from whole-genome sequencing, error rates that are low per-site can still yield millions of miscalled genotypes. Errors can be introduced anywhere along the long process, from sampling to genotyping. The frequency of error often depends on the sequencing technology employed. With next-generation sequencing (NGS), even reads that are perfectly mapped and free of base-calling errors can lead to miscalled diploid genotypes due to the random over-sampling of one allele during amplification. Genotyping errors can profoundly hinder genetic analyses, for instance by reducing power in linkage and association studies (Abecasis *et al.* 2001; Gordon *et al.* 2002; Ahn *et al.* 2007). Studies interested in identifying rare variants are especially sensitive to these errors, in the context of either disease (Powers *et al.* 2011; Yan *et al.* 2016) or the rate of *de novo* mutation (Ségurel *et al.* 2014; Carlson *et al.* 2018). The consequences of genotyping errors for biological conclusions can often be mitigated by explicitly including the possibility of errors in genetic analyses (Sobel *et al.* 2002; Cartwright *et al.* 2007; Lebrec *et al.* 2008).

Methods for attenuating the effects of error often require investigators to use an estimate of error rates (Pompanon *et al.* 2005). While error rates can broadly be classified by the type of sequencing or genotyping technology employed, each experiment will have a different error rate. Identical pipelines for generating genotype data with NGS can lead to rates of error that vary between different samples and cohorts (Dohm *et al.* 2008; Huang *et al.* 2009). For example, the population frequency of each variant in a cohort can be used to improve the accuracy of genotype calls, as in the GenotypeGVCFs workflow (Poplin *et al.* 2017). While this approach increases the confidence in each genotype call, genotype error rates can become dependent on the rarity of variants at a locus.

There are several different approaches for estimating genotyping error rates in a given experiment. The most straightforward is some form of replication, where sequencing or genotyping on one or more samples is repeated—or performed at higher read-coverage with NGS—and compared to the original results (e.g. Wall *et al.* 2014; Pfeiffer *et al.* 2018; Ma *et al.* 2019). Aside from the potential to be cost-prohibitive, this approach generates a rate of discordance between replicates rather than a true estimate of the error rate. Robust approaches to identifying genotyping errors and estimating their rates typically leverage pedigree information. These approaches identify errors by finding discordance between the observed genotypes and those expected from the laws of Mendelian inheritance (Douglas *et al.* 2002; Hao *et al.* 2004; Saunders *et al.* 2007; The 1000 Genomes Project Consortium 2015).

In this study, we develop a method to estimate the rates at which different types of genotyping errors occur. We focus on errors at biallelic sites and distinguish between errors that involve reference versus non-reference alleles. Our method is useful for establishing a distinct genotyping error rate for genetic analyses of rare variants, which are typically non-reference. We apply pedigree information to track the transmission of haplotypes and to identify errors by looking for sites that have genotypes that cannot be the result of a faithful transmission event. Loci that do not possess the expected phase within a haplotype block must be the result of either: a *de novo* mutation, a gene conversion event, or a genotyping error. Genotyping errors are by far the most common of these phenomena (Table 1). Detection of unlikely genotypes from haplotype phase has previously been used to successfully identify genotyping errors (Abecasis *et al.* 2002; Becker *et al.* 2006; O’Connell *et al.* 2014; Kothiyal *et al.* 2019). What we add here is an error model to explicitly determine the genotyping error rate, and the ability to distinguish rates for different types of errors.

**Table 1.** Approximate per-site rates for phenomena leading to phase inconsistency

|  |  |
| --- | --- |
| NGS genotyping error | $10^{-3}$ |
| gene conversion | $10^{-7}$ |
| recombination | $10^{-8}$ |
| *de novo* mutation | $10^{-8}$ |

We apply our new method to a dataset of genotypes collected from the whole-genome sequencing of a set of owl monkey (*Aotus nancymaae*) pedigrees (Thomas *et al.* 2018). Among sites that could be phased with pedigree information, we found a significant difference in the direction in which phase errors occurred. This departure forms the signal for estimating the rate of genotyping error. Estimated error rates were significantly different among the genotypes, with the most common error being a homozygous reference site miscalled as heterozygous. The principles of our method can be applied to determine the rate of different types of genotyping error in any dataset where phase errors can be identified.

## Materials and Methods

We develop a method to estimate genotyping error rates from whole-genome data by examining autosomal sites that can be unambiguously phased from a three-generation pedigree. At phase-informative sites, the transmission of haplotypes can be determined according to the Mendelian laws of inheritance. When the genotype of the child does not match the haplotypic combination transmitted by the parents, a genotyping error is the most likely cause. The frequency of such phase violations depends on the rate of genotyping error and the relevant site frequency in the sample. The expected frequencies of different phase violations can thus be compared to the observed frequencies to estimate error rates. We focused on building an error model for three-generation pedigrees along a single line of descent (Fig. 1a), though genotypes can be phased in two-generation pedigrees when there are more than two siblings (Fledel-Alon *et al.* 2009; Roach *et al.* 2011). We also restricted ourselves to biallelic sites, the most common form of variation across the genome. The six possible miscalls at a biallelic site are labeled in Table 2.

**Table 2.**
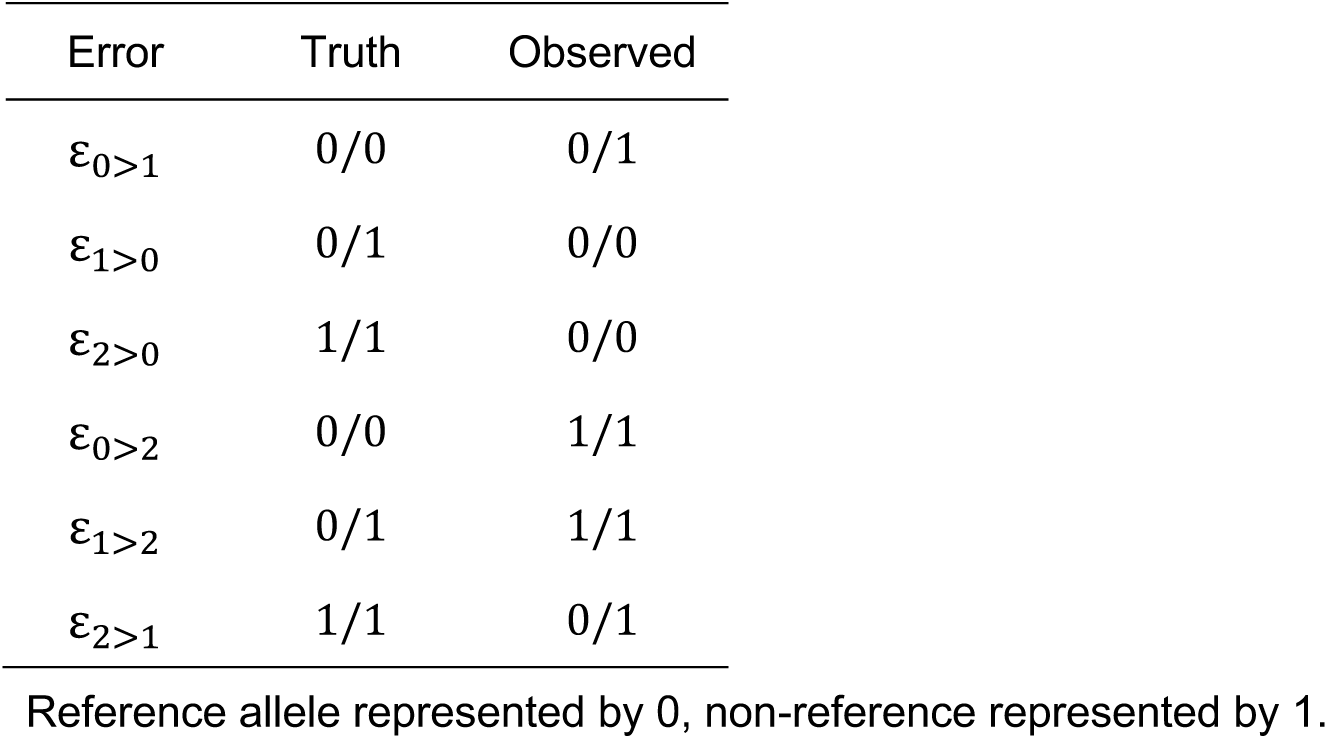
Genotyping errors at a biallelic site

**Figure 1.**
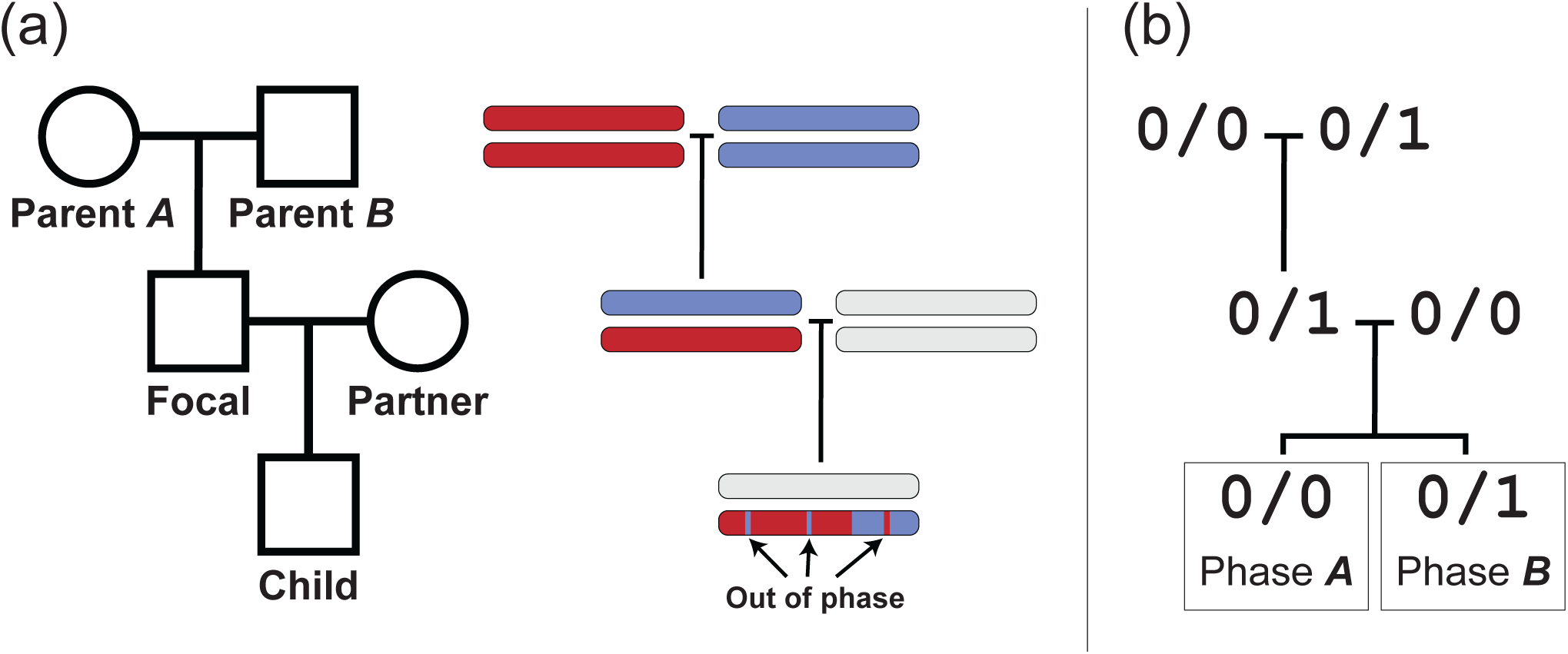
Pedigree and phase in a three-generation pedigree along a single line of descent. (a) The pedigree of five individuals and a diagram of the haplotype traced along three generations. Sites transmitted to the child can be out of phase within a haplotype block. (b) Genotypes at an example phase-informative site. The phase of the child genotype can be traced to either Parent A or Parent B.

### Expected number of phase violations

At phase-informative sites, the focal individual in a three-generation pedigree produces gametes with traceable phase. Within a block without crossovers, the haplotype inherited from the focal individual can be identified as being derived from either the maternally or paternally inherited chromosome of the focal individual. With a three-generation pedigree (Fig 1a), phase-informative sites are those where: the focal individual is heterozygous, the parents of the focal individual are not both heterozygous, and the breeding partner of the focal individual and the child are not both heterozygous.

A genotyping error in any of the five individuals from the pedigree can cause an informative site to appear out-of-phase with its neighbors. For example, for a set of genotypes as in Figure 1b, a child called as heterozygous at a site in a parent *A* haplotype block could be an ε_0>1_ error, a miscalled homozygous child. Similarly, a homozygous child at a site in a parent *B* block could be an ε_1>0_ error. A miscalled child genotype is the most straightforward cause of a phase violation, but errors in other individuals also create apparent phase violations. Using again the genotypic combination in Figure 1b—if parent *A* is miscalled as homozygous alternate instead of homozygous reference (an ε_2>0_ error; Table 2) a site in a parent *A* block would be expected to carry the alternate allele. When parent *A* is miscalled in this way, the child will be heterozygous for a site in a parent *A* block and would appear to be a phase violation.

The expected number of violations is the sum of potential phase violations from miscalls in all individuals across the pedigree. We estimate the number of violations for each miscall as a product of the respective genotypic combination frequencies across the genome and the corresponding error rate. We demonstrate this rationale below for the genotypic combination matching the example in Figure 1b, calculating the expected number of such phase-violating sites from phase *A* haplotype blocks. In this calculation, we assumed that out-of-phase informative sites were the result of no more than one genotyping error among the sampled individuals for a given site.

Genotypic combinations are abbreviated as *G*_*x*_, where *x* is composed of individual genotypes represented by a single digit: 0, 1, and 2 for homozygous reference, heterozygous, and homozygous alternate respectively. These are ordered in the subscript as: parent *A*, parent *B*, focal individual, partner of focal, and child. Let *n*_*x*_ be the number of such genotypic combinations and ε_i>j_ be the genotyping error rate as in Table 2. The genotypic combinations and possible miscalls leading to a phase violation for a site like the one depicted in Figure 1b are then:

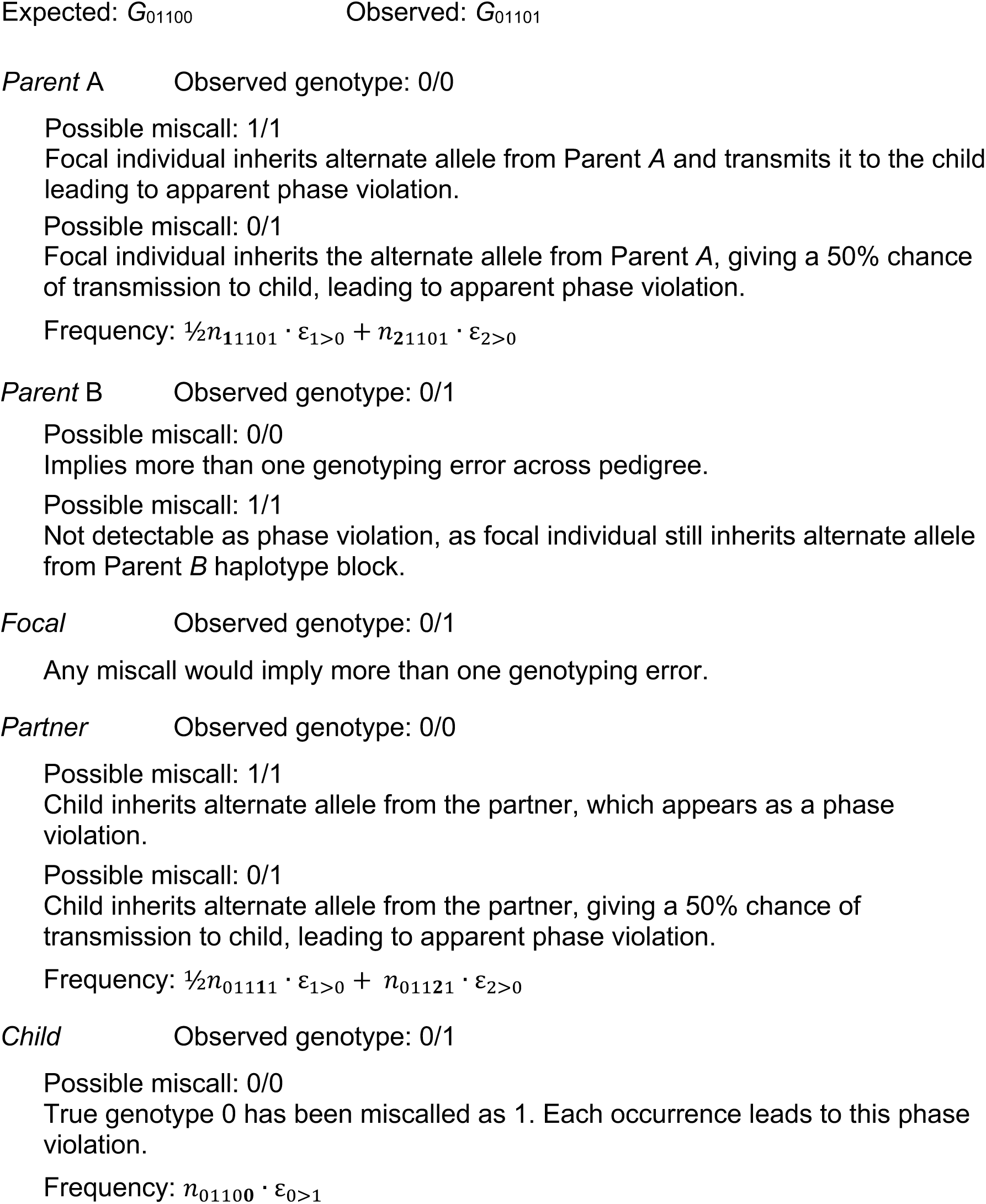

The expected number of phase violations for this genotypic combination can be written as

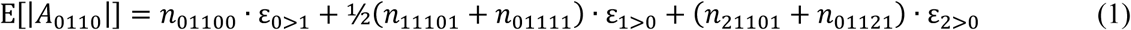

where *A*_y_ is a phase violation in the parent *A* haplotype block, with genotypic combination *y* and |*A*_y_| is the number of such violations across the genome. Here, *y* is the abbreviated phase-informative genotypic combination, listing abbreviated genotypes for the first four individuals as ordered in *x* for *G*_x_.

In pedigrees with five individuals there are 18 genotypic combinations that are phase-informative. Table 3 lists each of these combinations and the child genotypes that indicate a phase violation. Violations at sites with the haplotype from Parent *A* and Parent *B* are labeled as *A*_y_ and *B*_y_ respectively. Each genotypic combination can be evaluated to determine the total number of expected phase violations. Several symmetries between genotypic combinations reduce the number of unique calculations. For example, each violation has a matching pair where the phase and the genotypes of Parent *A* and Parent *B* are swapped. The frequency of each of these two classes is expected to be equal across the genome due to independent assortment. The matching violation to *A*_0110_ in the example above is *B*_1010_, and the expected frequency can be calculated by swapping the two parental genotypes in each term as

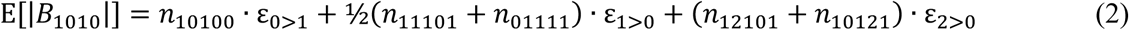

**Table 3.** Detectable phase violations in a three-generation pedigree

| Parent<br>A | Parent<br>B | Focal | Partner | Phase<br>Violation | Child<br>(Expected) | Child<br>(Observed) | Phase<br>Violation | Child<br>(Expected) | Child<br>(Observed) |
| --- | --- | --- | --- | --- | --- | --- | --- | --- | --- |
| 0/0 | 0/1 | 0/1 | 0/0 | $A_{0110}$ | 0/0 | 0/1 | $B_{0110}$ | 0/1 | 0/0 |
| 0/0 | 0/1 | 0/1 | 0/1 | $A_{0111}$ | 0/0; 0/1 | 1/1 | $B_{0111}$ | 1/1; 0/1 | 0/0 |
| 0/0 | 0/1 | 0/1 | 1/1 | $A_{0112}$ | 0/1 | 1/1 | $B_{0112}$ | 1/1 | 0/1 |
| 1/1 | 0/1 | 0/1 | 0/0 | $A_{2110}$ | 0/1 | 0/0 | $B_{2110}$ | 0/0 | 0/1 |
| 1/1 | 0/1 | 0/1 | 0/1 | $A_{2111}$ | 1/1; 0/1 | 0/0 | $B_{2111}$ | 0/0; 0/1 | 1/1 |
| 1/1 | 0/1 | 0/1 | 1/1 | $A_{2112}$ | 1/1 | 0/1 | $B_{2112}$ | 0/1 | 1/1 |
| 0/0 | 1/1 | 0/1 | 0/0 | $A_{0210}$ | 0/0 | 0/1 | $B_{0210}$ | 0/1 | 0/0 |
| 0/0 | 1/1 | 0/1 | 0/1 | $A_{0211}$ | 0/0; 0/1 | 1/1 | $B_{0211}$ | 1/1; 0/1 | 0/0 |
| 0/0 | 1/1 | 0/1 | 1/1 | $A_{0212}$ | 0/1 | 1/1 | $B_{0212}$ | 1/1 | 0/1 |
| 1/1 | 0/0 | 0/1 | 1/1 | $A_{2012}$ | 1/1 | 0/1 | $B_{2012}$ | 0/1 | 1/1 |
| 1/1 | 0/0 | 0/1 | 0/1 | $A_{2011}$ | 1/1; 0/1 | 0/0 | $B_{2011}$ | 0/0; 0/1 | 1/1 |
| 1/1 | 0/0 | 0/1 | 0/0 | $A_{2010}$ | 0/1 | 0/0 | $B_{2010}$ | 0/0 | 0/1 |
| 0/1 | 1/1 | 0/1 | 1/1 | $A_{1212}$ | 0/1 | 1/1 | $B_{1212}$ | 1/1 | 0/1 |
| 0/1 | 1/1 | 0/1 | 0/1 | $A_{1211}$ | 0/0; 0/1 | 1/1 | $B_{1211}$ | 1/1; 0/1 | 0/0 |
| 0/1 | 1/1 | 0/1 | 0/0 | $A_{1210}$ | 0/0 | 0/1 | $B_{1210}$ | 0/1 | 0/0 |
| 0/1 | 0/0 | 0/1 | 1/1 | $A_{1012}$ | 1/1 | 0/1 | $B_{1012}$ | 0/1 | 1/1 |
| 0/1 | 0/0 | 0/1 | 0/1 | $A_{1011}$ | 1/1; 0/1 | 0/0 | $B_{1011}$ | 0/0; 0/1 | 1/1 |
| 0/1 | 0/0 | 0/1 | 0/0 | $A_{1010}$ | 0/1 | 0/0 | $B_{1010}$ | 0/0 | 0/1 |

Reference allele represented by 0, non-reference allele represented by 1.
Violations are mirrored in the second half of the table (Parent A and Parent B genotypes swapped, A and B violation swapped).

Similarly, each violation can be paired with a case in which homozygous reference and homozygous alternate genotypes are swapped. The expected frequency of violations has the same form once these genotypes and error rates are swapped. For *A*_0110_, the matching violation is *A*_2112_ with expected frequency

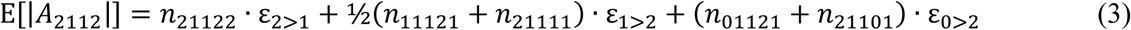

Derivations of the expected frequencies for the remaining set of phase violations can be found in the Supplemental Material (Appendix S1).

### Estimating the error rates

Taken together, the expected number of phase violations from the genotypic combinations in Table 3 form a linear system of equations that can be represented by a matrix equation. Let **V**_**A**_ be a vector for the number of observed sites that violate haplotype block *A* across the genome, as ordered by row in Table 3; that is, **V**_**A**_ = ⟨|*A*_0110_|, |*A*_0111_|, |*A*_0112_|, …, |*A*_1011_|, |*A*_1010_|⟩. The number of observed sites that violate block *B* is represented by **V**_**B**_, again as ordered in Table 3. Let 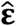 be a vector of estimators for each type of error rate, 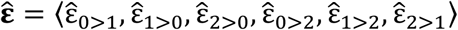. The error model can then be represented as

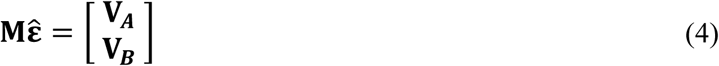

where the matrix **M** contains the coefficients of the linear equations from the expected frequencies of genotyping errors for each violation class. We can divide **M** into two submatrices based on the coefficients for violations in phase *A* and phase *B* as

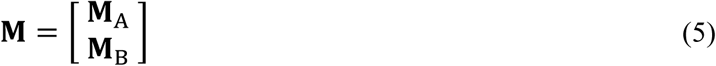

Rows in **M**_**A**_ and **M**_**B**_ are identical for violations in phase *A* and phase *B* that share the same expected frequencies (e.g. *A*_0110_ and *B*_1010_, described in the previous section). The violations in Table 3 are ordered so that **M**_**A**_ and **M**_**B**_ can be related by an exchange matrix as

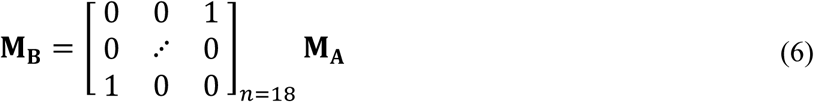

Coefficients from equations for each of the respective expected phase *A* violations (see Supplemental Information) form the **M**_**A**_ matrix:

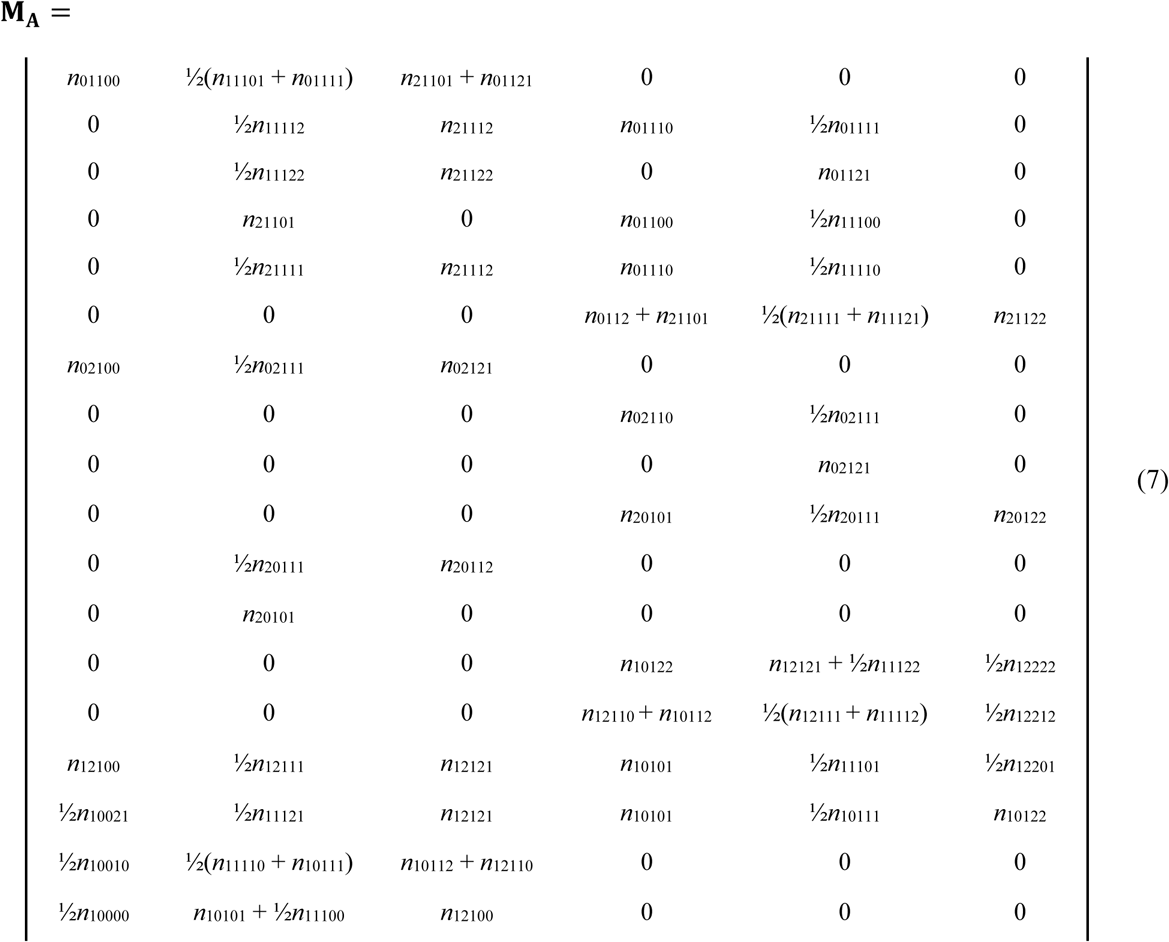

Equation 4 is an overdetermined linear system—there are many more equations than rates to be estimated. We can fit the model using a linear least squares approach, solving for 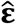 by taking

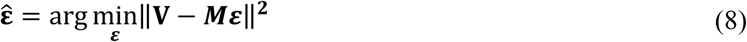

Our implementation takes as initial input genotypes in the Variant Call Format (vcf) and phases for haplotype blocks in Browser Extensible Data (BED) format. We fed this data in abbreviated form into R (v. 3.5.0) and solved for the error rate estimators in Equation 8 with the linear optimization algorithm L-BFGS-B as implemented in the base stats package. The Python (v. 3.6.1) script used to abbreviate the genotype-phase combinations, and the R script applying the algorithm, are available on GitHub.

### Simulating phase violations

We tested the performance of our method on simulated genotype combinations at biallelic sites from pedigrees as in Fig. 1a with simulated errors. The phase at each site was randomly assigned, consistent with the rules of Mendelian inheritance, and individual genotypes each had a chance of being in error. To quickly simulate a large number of such genotypes and errors, we drew from a probability distribution of genotypic combinations instead of individual genotypes. We assumed a neutral site frequency spectrum for each site and that the parents and partner of the focal individual were all unrelated to each other. Assuming that the three unrelated individuals in the pedigree are from a subset of *N* (minimum *N* = 3) unrelated genotyped individuals that have *S* segregating sites, the probability of a given unrelated genotypic combination, *U*_*x*_, can be written as

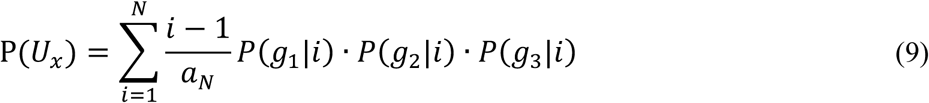

where *a*_N_ is the Watterson correction factor

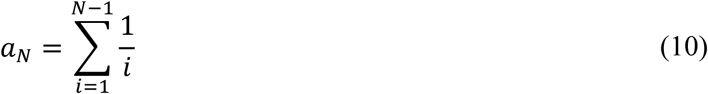

*g*_*n*_ is the genotype of the *n*th-individual in the combination, and the conditional genotype frequency is given by

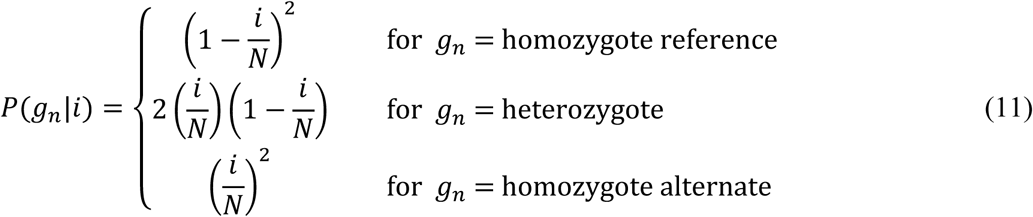

The count for the 27 possible genotypic combinations, ⟨*C*_1_, …, *C*_27_⟩, can then be drawn from a multinomial with parameters ⟨P(*U*_1_), …, P(*U*_27_)⟩ for the number of segregating sites, *S*. For each genotypic combination, the *C*_*x*_ outcomes were divided among the possible genotypes of the focal individual, based on the parental genotypes in *U*_*x*_. These counts were apportioned by drawing from a multinomial with an equal probability for each of the four possible phase and genotype combinations. These branching outcomes were in turn further divided by the possible child genotype from the focal and partner genotypes, drawn again from a multinomial with equal probability for each of the four possible outcomes. Counts across all branches were then summed to give a total count, *n*_*x,p*_, for each unique genotype and phase combination, *x* and *p*.

We added errors to these counts by iterating over the genotypes in each combination, *x*, and drawing errors for the *n*_*x,p*_ sites. Let the genotype be a vector, *g* = {⟨1,0,0⟩, ⟨0,1,0⟩, ⟨0,0,1⟩}, for the homozygous reference, heterozygous, and homozygous alternate genotypes respectively. Given a set of error probabilities, *ε*_*i*>*j*_, for each type of error as listed in Table 2, the probability of a genotypic transition can be written as *g* · **E**, where

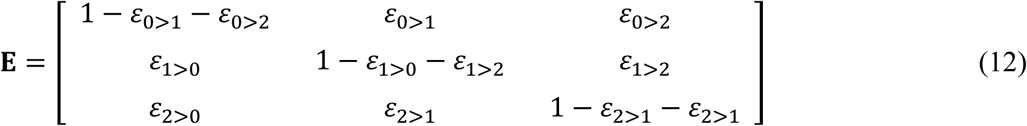

For each genotype at each combination, the *n*_*x*_,_*p*_ counts are divided by drawing from a multinomial using the probability for genotypic transition while retaining the phase. Unique genotype and phase combinations across all branches are then summed for the final counts.

Simulations following the above strategy were implemented in Python (v. 3.6.1) with the NumPy package (v. 1.13.3).

### Data Availability

Raw sequence data for the owl monkey dataset is available from NCBI BioProject: PRJNA451475. Abbreviated counts of genotype-phase combinations from the owl monkey dataset are available at FigShare. Code used in analyses and simulations are publicly available on GitHub (https://github.com/Wang-RJ/genotypeErrors).

## Results

### Accuracy of error estimators in simulations

We simulated genotypes with varying error rates and numbers of segregating sites, maintaining the sample size of unrelated genotyped individuals at *N* = 20. Simulated error rates were drawn from two distributions, a log-uniform distribution (range: [10^−4^, 10^−2^]) and a log-normal distribution (μ = 10^−3^, σ = 0.5). We simulated 1000 genotyped pedigrees of 5 individuals for each combination of parameters tested. In the absence of genotyping errors, the frequency of parental phases at informative sites is expected to be equal due to independent assortment. However, genotyping errors cause apparent phase violations to occur at different frequencies based on the genotypic combination of individuals in the pedigree. Figure 2a shows an example of how the parental phase at detectable violations consistently varies from the balanced phases at informative sites. The frequency of simulated genotypic combinations also reflects the neutral site frequency spectrum (see Materials and Methods).

**Figure 2.**
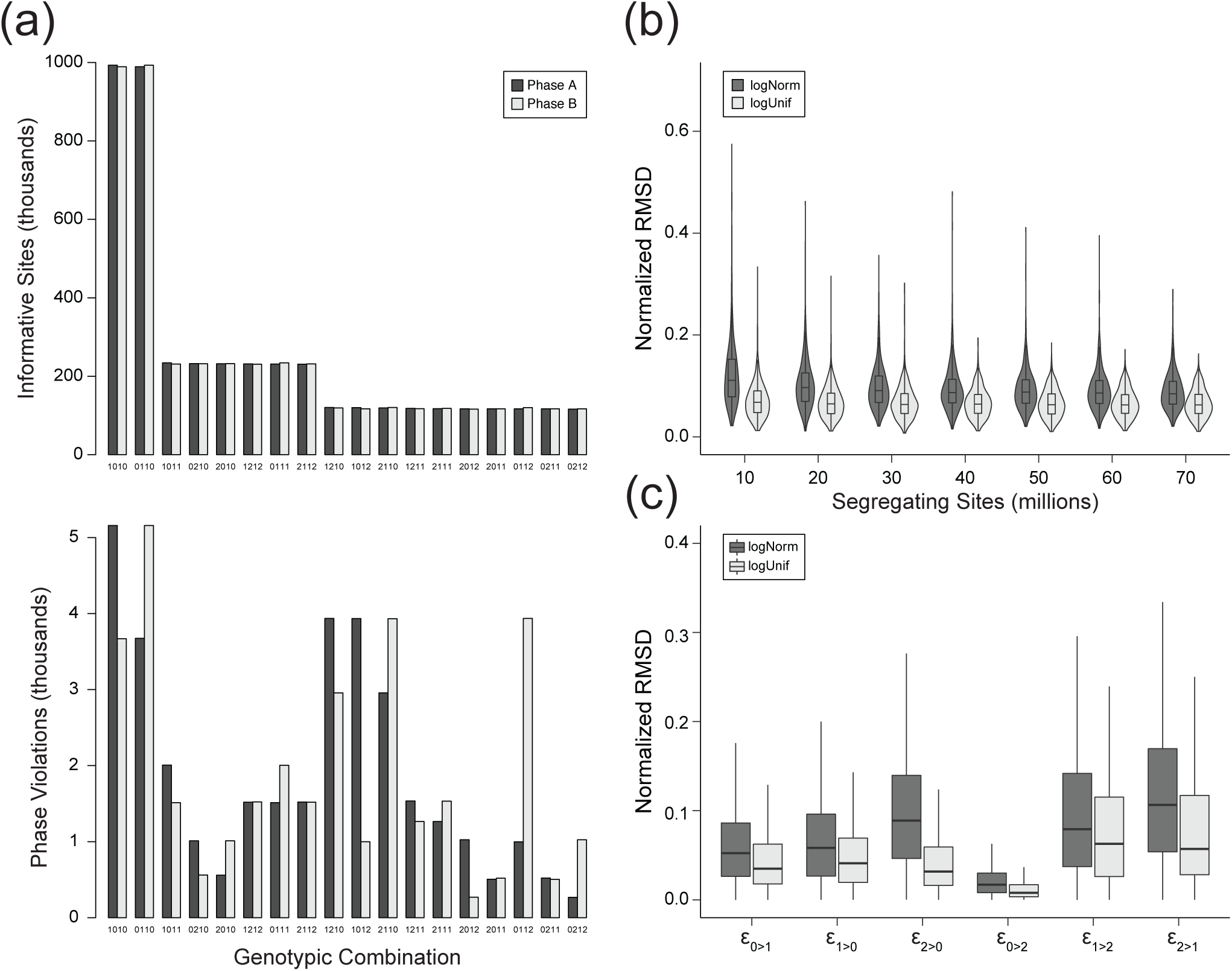
Estimating simulated genotyping error rates from phase violations. (a) Typical genotypic combinations from simulations, ordered by the expected frequency of informative sites. Phases are expected to occur at equal frequency for informative sites (top), but violations due to genotyping error do not occur equally (bottom). Depicted are the mean frequencies from 1,000 simulations with error rates drawn from a log-uniform distribution. (b) Total deviation of rate estimators decreases modestly with more segregating sites in simulated samples. (c) Deviation of individual estimators from actual rates used in simulations.

We assessed the accuracy of our error rate estimation in these simulations by calculating the normalized root mean square deviation (NRMSD). The deviation between the estimators and the simulated rate was normalized by the range of error rates as

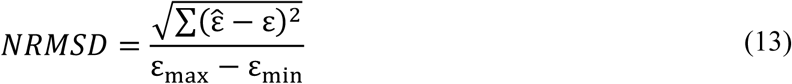

where ε_max_ and ε_min_ are the maximum and minimum error rates in each simulation.

There was little change to the estimators’ normalized deviation with an increasing number of segregating sites for simulated error rates drawn from either the log-normal or log-uniform distribution. The mean NRMSD for simulated rates drawn from the log-normal distribution was slightly higher, ranging from 9.0% to 12.2%, than from rates drawn from the log-uniform distribution, from 6.5% to 7.1%, for simulations with between 10 and 70 million segregating sites (Fig. 2b). These results indicate that our method of estimating error rates is robust across populations with different levels of nucleotide diversity.

We also assessed our method’s accuracy for each estimator, normalizing the RMSD to the median difference between maximum and minimum error rates across all simulations (Fig. 2c). We found 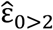 to be the most accurately estimated, with a mean NRMSD of 2.1% and 1.2%, for simulated errors drawn from the log-normal and log-uniform respectively. The estimators 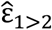 and 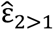 were the least accurate, with mean NRMSDs ranging from 8.0% to 12.4%. The accuracy of the estimator for error rates at the most frequent genotype, 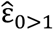, was intermediate with an NRMSD of 4.3% and 6.1% for respective draws from the log-normal and log-uniform distributions.

### Application to phased owl monkey genotypes

We applied the method developed here to a dataset of genotypes from the whole-genome sequencing of owl monkey (*Aotus nancymaae*). Individuals in this dataset were part of several three-generation pedigrees, allowing us to unambiguously phase focal individuals (as in Fig. 1). We selected four unrelated pedigrees with genotypes from twenty total individuals for our analysis. These samples were sequenced to approximately 35× coverage on an Illumina HiSeq-X with 150 bp paired-end reads. Single nucleotide polymorphisms (SNPs) in this dataset were called with GATK (version 3.3.0) following best practices (Van der Auwera *et al.* 2013; Poplin *et al.* 2017), and genotypes were phased with PhaseByTransmission (Francioli *et al.* 2017) and assembled into haplotype blocks under the assumption that there would be at most one recombination crossover per Mb interval (Venn *et al.* 2014; Smeds *et al.* 2016). Complete details on the methods used to generate this dataset are available in Thomas *et al.* (2018).

We selected genotypes from biallelic SNPs on autosomes for the subsequent analyses. Based on genotypic combinations, we were able to identify approximately 5.5 million phase-informative sites in each of the pedigrees. These sites were on haplotype blocks that covered, on average, 2.46 Gb out of the 2.86 Gb genome. Among the informative sites, an average of 25,007 were out-of-phase in each family. Frequencies for the different genotypic combinations showed a similar pattern to those seen in simulations: a balanced set of phases among all informative sites and an unequal set of phases among the violations (Fig. S1). Note that the proportion of out-of-phase sites relative to the total number of informative sites does not represent an error rate because potential miscalls are spread across the genotypes of multiple individuals in the pedigree.

Using the frequencies of different phase violations, we estimated the genotyping error rates for each family with Equation 4. Estimates from each of the four pedigrees are depicted in Figure 3. The highest rates of error appear to be from distinguishing between homozygous reference and heterozygous genotypes, with mean rates ε_0>1_ = 2.9 × 10^−3^ and ε_1>0_ = 2.9 × 10^−3^ per bp. The rate at which homozygotes for the alternate allele were miscalled as heterozygotes was also appreciable, mean rate ε_2>1_ = 2.1 × 10^−3^ per bp. In contrast, our estimate of the rate at which homozygous reference genotypes were mistaken as homozygous alternate was zero in all four pedigrees.

**Figure 3.**
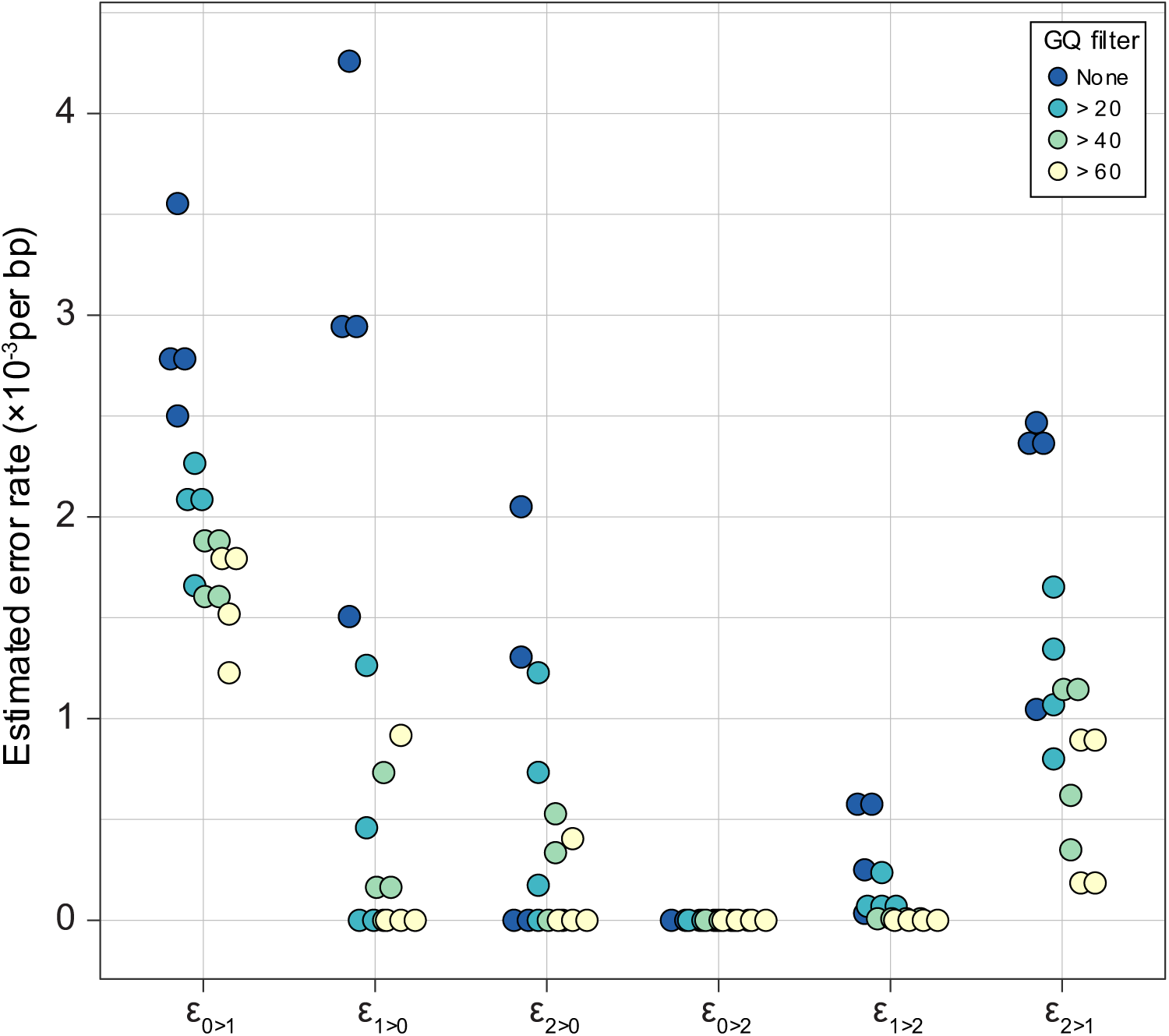
Error rates estimated from sequencing four pedigrees of owl monkeys. Each point represents an estimate from one pedigree with genotypes filtered as described in the legend. Estimated genotyping error rates consistently decline with the stringency of the GQ filter.

We repeated the analysis with genotypes that were filtered by genotype quality (GQ) as calculated in GATK. The value of GQ is based on the difference between the probability of the called genotype and the next most likely genotype. Genotypes associated with higher GQ scores are typically interpreted as being more accurate. As expected, the estimated error rates decreased as we removed sites with more stringent GQ filters (Fig. 3; Table S1). Filtering by GQ appears to be more effective at removing certain types of errors than others, with the most dramatic reduction in heterozygote false negatives, that is ε_1>0_.

We also calculated an average error rate by weighting each estimator with the genome-wide frequency of the corresponding genotype (e.g. the frequency of homozygous reference genotypes for ε_0>1_). As with individual estimators, increasing stringency of the GQ filter reduced the overall error rate (Table S1). We estimated an overall error rate of 3.0 × 10^−3^ per bp, which was reduced to 1.1 × 10^−3^ per bp when genotypes were required to have a GQ > 60.

## Discussion

We have developed a method to estimate genotyping error rates for different types of errors at biallelic loci. Leveraging pedigree information, our method directly estimates underlying error rates, rather than the discordance between experiments obtained by other approaches. Our method is more robust than those that consider only Mendelian violations in a trio of individuals because of additional transmission information, reliance on multiple biological phenomena, and the ability to distinguish different types of errors.

Our estimate of the overall genotyping error rate in the owl monkey samples is comparable to estimates calculated from discordance between replicate sequencing experiments. Wall et al. (2014) inferred a genotyping error rate of 1s.18 × 10^−3^ per bp on the Illumina HiSeq platform at GQ >= 40, remarkably similar to our overall estimate of 1.3 × 10^−3^ per bp at the same level of filtering stringency (Table S1). Though their resequencing approach does not distinguish between all types of miscalls, they report a false positive rate for heterozygotes (ε_0>1_) that is much higher than the false negative rate (ε_1>0_), consistent with our findings after the application of filters on GQ.

Heterozygote false positives at homozygous reference sites occur at the highest rate among all errors, after modest GQ filtering. As the most common type of site in the genome, they are also the most common genotyping error. That the lowest error rates were for erroneously called non-reference homozygotes may have been expected. The relative rarity of the non-reference allele leads to caution in calling non-reference homozygotes by most genotyping methods. More surprising is the uneven effect of filtering by GQ across error types. Heterozygote false negatives, in particular, were dramatically reduced by this filter. If the disparity between heterozygote false positive and false negative rates is common across NGS experiments, studies that seek rare variants may not be calibrated appropriately when assuming a single error rate. While genotyping errors for low-frequency variants may have little effect for many analyses, studies looking to identify *de novo* mutations in order to estimate mutation rates are very sensitive to miscalls of these variants.

The effect of genotype filters on the false negative rate can be difficult to quantify; false positives, on the other hand, can be detected by confirming candidate sites with, for example, Sanger sequencing. We demonstrated that our method allows for an analytic estimate of the false negative rate with varying degrees of genotype filtering. At first glance, the order of magnitude difference in heterozygote false negative rates when filtering by GQ might suggest a corresponding difference in the number of false negatives in studies of *de novo* mutation. These studies, however, generally employ additional downstream filters to strictly control for the high number of false positives (Ramu *et al.* 2013; Wei *et al.* 2015). An estimate of each filter’s effect on the *de novo* false positive rate may be possible by applying our method to filtered genotypes, as we have done with GQ.

Simulations demonstrated the power of our approach to estimate error rates even in samples with low levels of diversity. They also indicated differences in the accuracy of the error estimators, but these were not pronounced for the most common types of sites. One caveat to our simulations is that we did not simulate entire haplotype blocks or inaccuracies in phase calling. Low levels of diversity and high rates of recombination can make accurate phasing more difficult, while low rates of recombination could result in an imbalance in the proportion of phases across the genome. The assumption of at most one genotyping error among samples per site may slightly inflate our estimates of the error rate. The potential for two or more genotyping errors is small (on the order of 10^−6^), and had a negligible effect in our simulations, but may be higher at sites prone to sequencing or assembly errors. Similarly, we ignored the effects of gene conversion and *de novo* mutation, as they are expected to occur at negligible rates compared to genotyping error (Table 1). Furthermore, the signal from a genotyping error and a gene conversion event are nearly identical, though careful filtering and selection of sites have been successful in identifying gene conversion events (e.g. Williams *et al.* 2015; Miller *et al.* 2016). Finally, we note that the estimated error rates are for genotyping from a set of called variants. The heterozygote false positive rate, ε_0>1_, for example, does not apply to invariant reference sites.

As genomic data continues to accumulate, the consideration of genotyping errors will remain an essential part of genetic analyses. Though we have focused mainly on whole-genome sequence data, our approach is generally applicable to any collection of genotype data (e.g. SNP-chips or exome sequencing) from pedigreed samples. Interest in sequencing individuals from families, as in studies seeking to identify *de novo* mutations (Goldmann *et al.* 2016; Thomas *et al.* 2018; Sasani *et al.* 2019), provide special opportunities for this method to be useful. Studies estimating *de novo* mutation rates may be particularly interested in distinguishing the rates for different types of genotyping errors. Differences in error rates will directly affect estimates of the false positive and false negative rates, and subsequent calculations of the mutation rate (Besenbacher *et al.* 2015; Kim *et al.* 2019). We have shown here that different types of errors indeed occur at different rates, necessitating their inclusion in such studies.

## Acknowledgements

We thank Gregg Thomas, Jeff Rogers, and Alan Harris for help with the owl monkey data, and Rafael Guerrero and Mark Hibbins for thoughtful discussions that led to the development of this manuscript. This work was funded by the Precision Health Initiative of Indiana University.

## Supplemental Material

**Figure S1.**
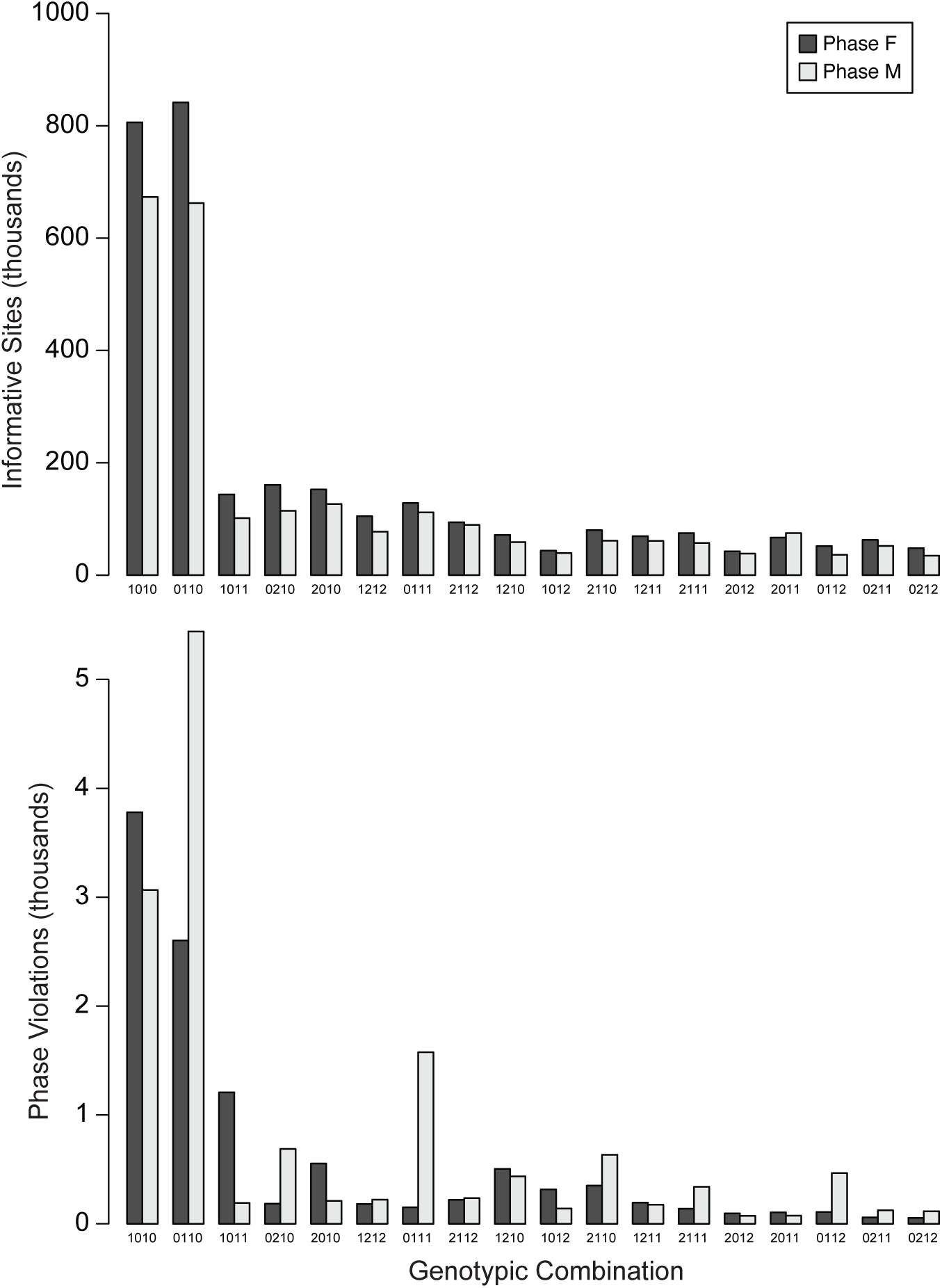
Mean number of informative sites and phase violations in the owl monkey dataset. Phases are at roughly equal frequencies for all informative sites relative to what is observed at violations. The overall frequency of genotypic combinations depends on the site frequency spectrum across the genome in owl monkeys. Genotypic combinations ordered as in Figure 2.

**Table S1.** Mean genotyping error rates (×10^−3^ per bp) estimated in the owl monkey dataset

| GQ filter | $\epsilon_{0>1}$ | $\epsilon_{1>0}$ | $\epsilon_{2>0}$ | $\epsilon_{0>2}$ | $\epsilon_{1>2}$ | $\epsilon_{2>1}$ | Genome-wide<sup>a</sup> |
| --- | --- | --- | --- | --- | --- | --- | --- |
| None | 2.9 (0.2) | 2.9 (0.6) | 0.8 (0.5) | 0.0 (0.0) | 0.4 (0.1) | 2.1 (0.3) | 3.0 (0.3) |
| > 20 | 2.0 (0.1) | 0.4 (0.3) | 0.5 (0.3) | 0.0 (0.0) | 0.1 (0.1) | 1.2 (0.2) | 1.6 (0.2) |
| > 40 | 1.7 (0.1) | 0.3 (0.2) | 0.2 (0.1) | 0.0 (0.0) | 0.0 (0.0) | 0.8 (0.2) | 1.3 (0.1) |
| > 60 | 1.6 (0.1) | 0.2 (0.2) | 0.1 (0.1) | 0.0 (0.0) | 0.0 (0.0) | 0.5 (0.2) | 1.1 (0.1) |

Standard errors in parentheses.
<sup>a</sup> Genome-wide estimated calculated by weighting rates with the normalized frequency of homozygote reference, heterozygote, and homozygote alternate genotypes in the owl monkey genome, respectively: 0.64, 0.25, and 0.11

## Appendix S1. Derivation of phase violation expected frequencies

Expected number of phase violations *A*_0111_, *A*_2111_

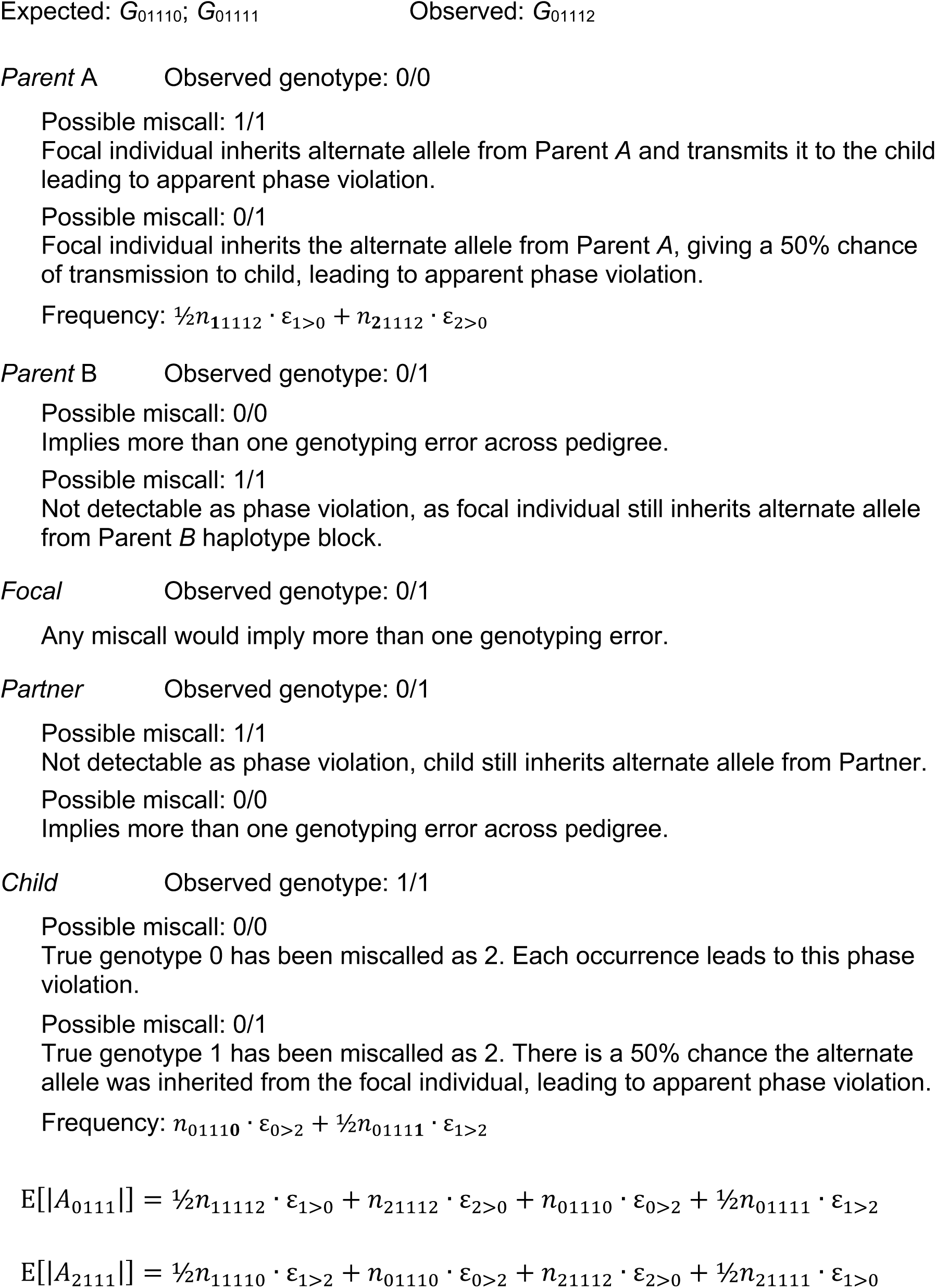

Expected number of phase violations *A*_0112_, *A*_2110_

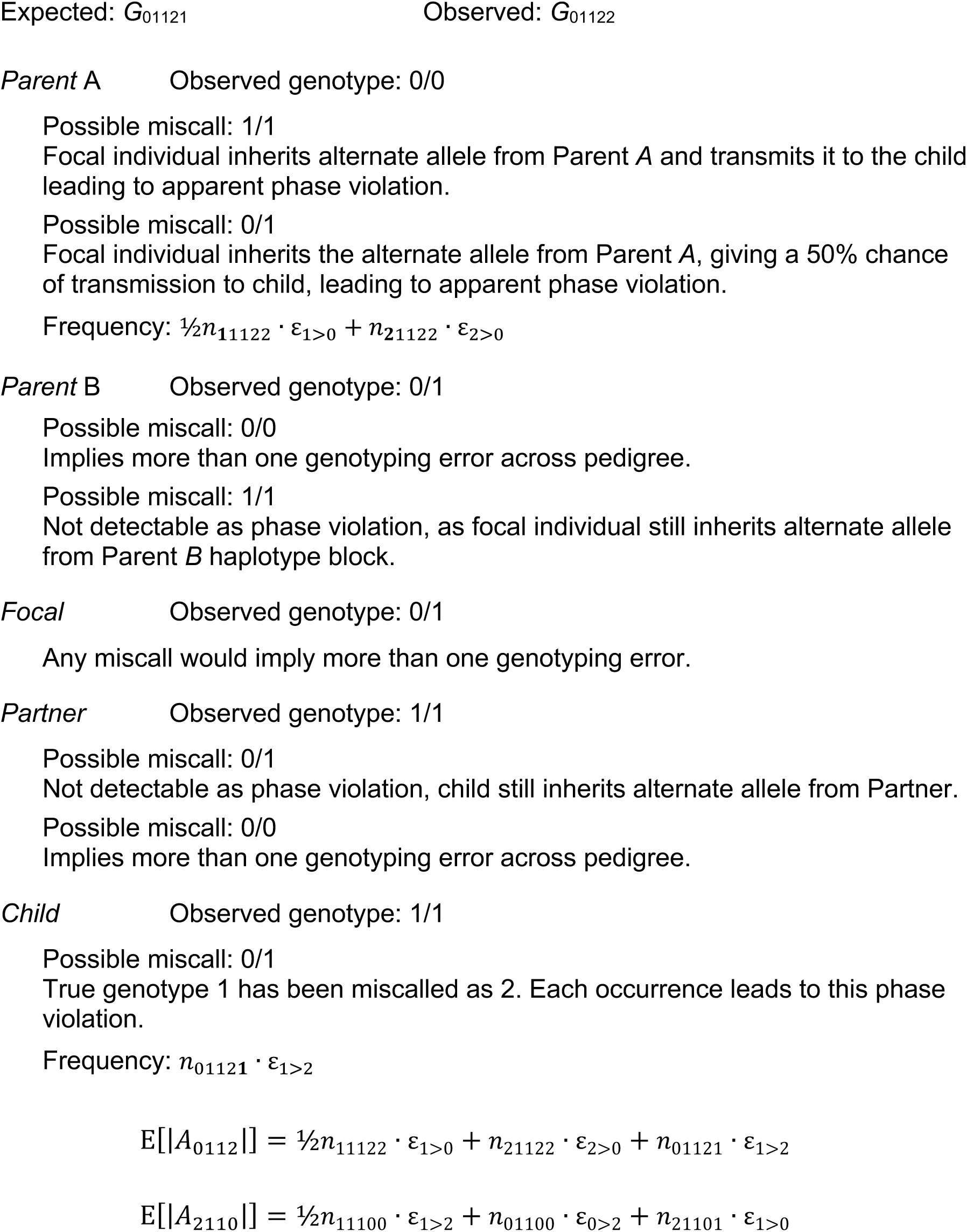

*Expected number of phase violations A*_0210_, *A*_2012_

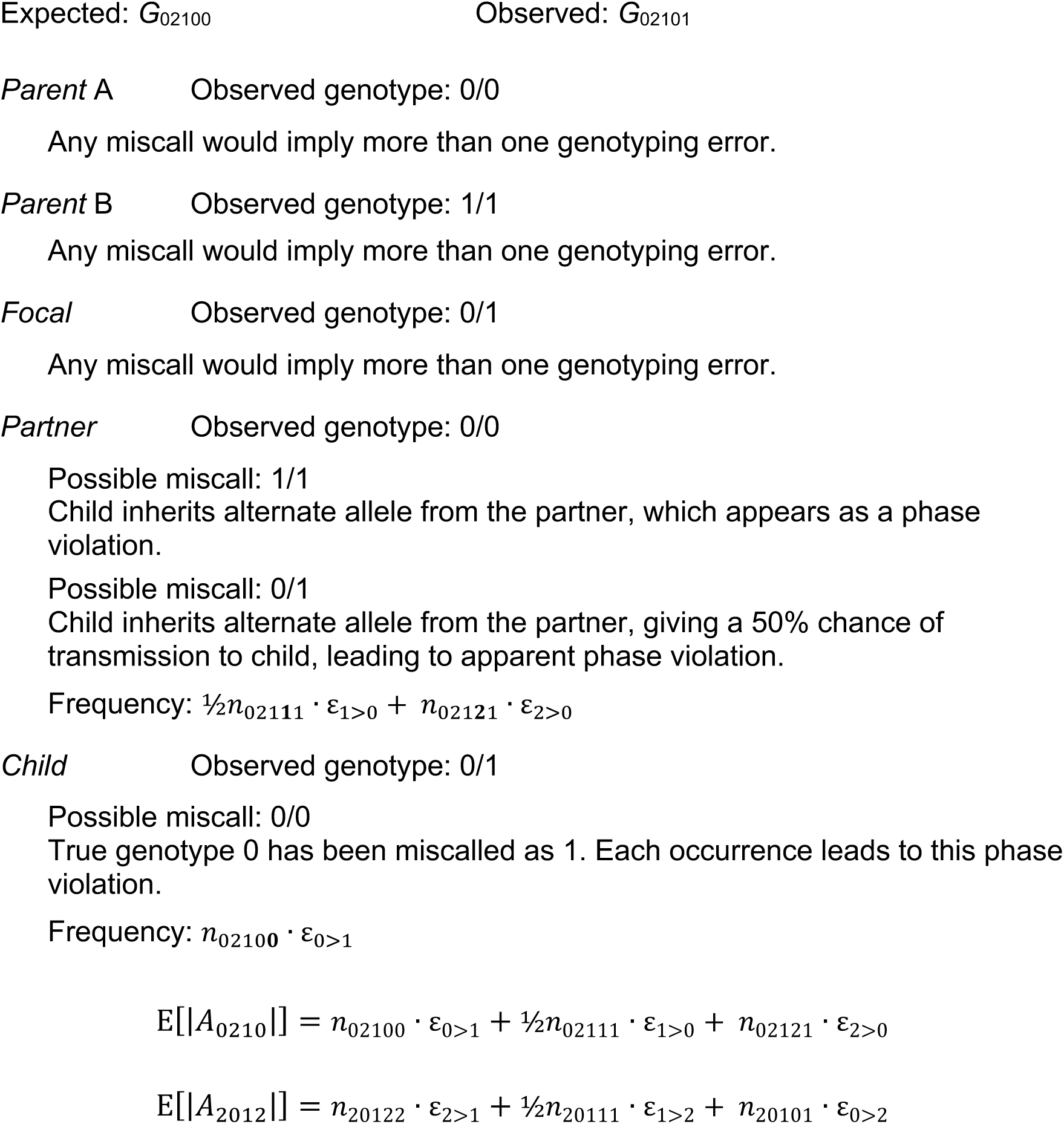

*Expected number of phase violations A*_0211_, *A*_2011_

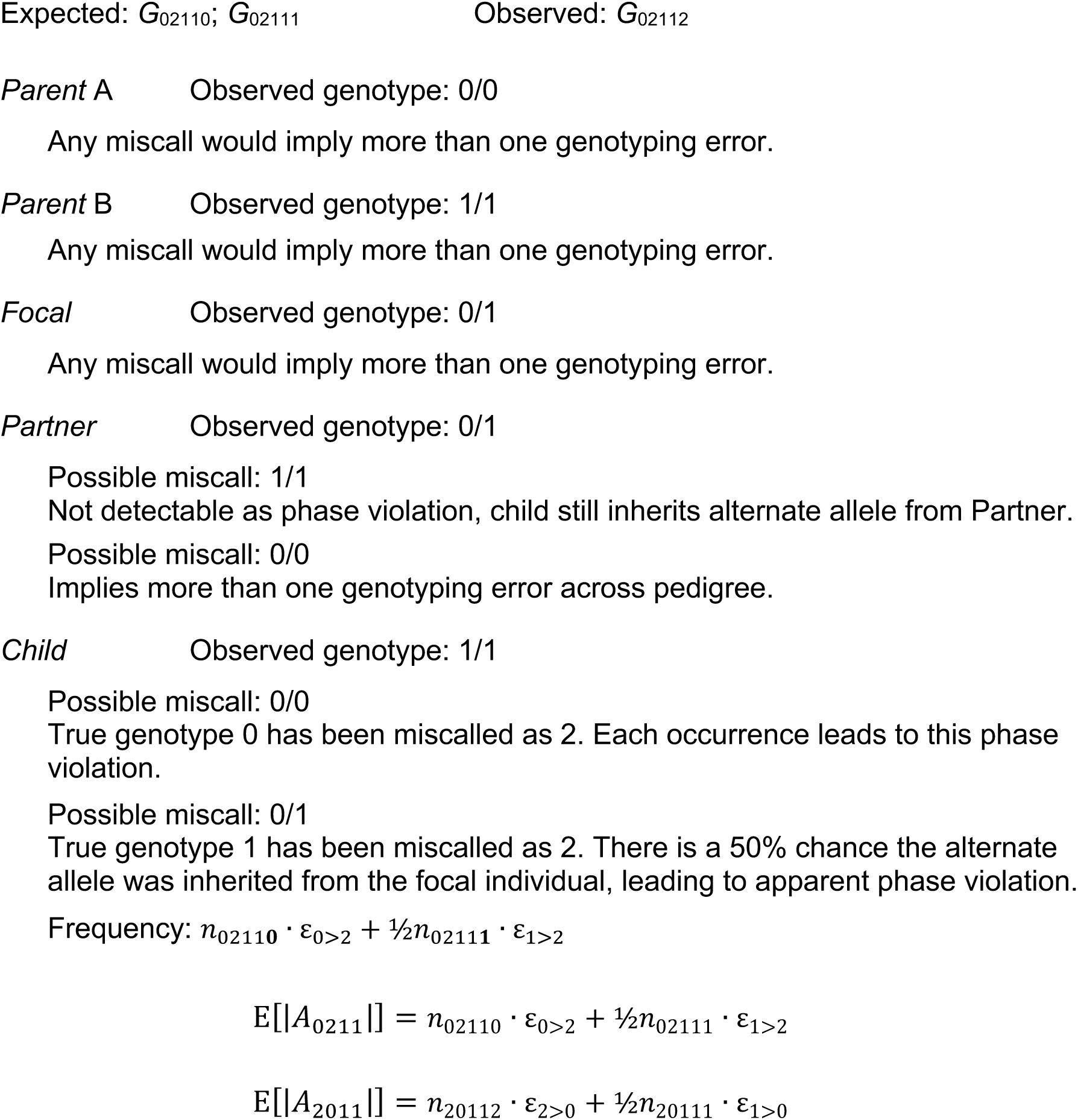

*Expected number of phase violations A*_0212_, *A*_2010_

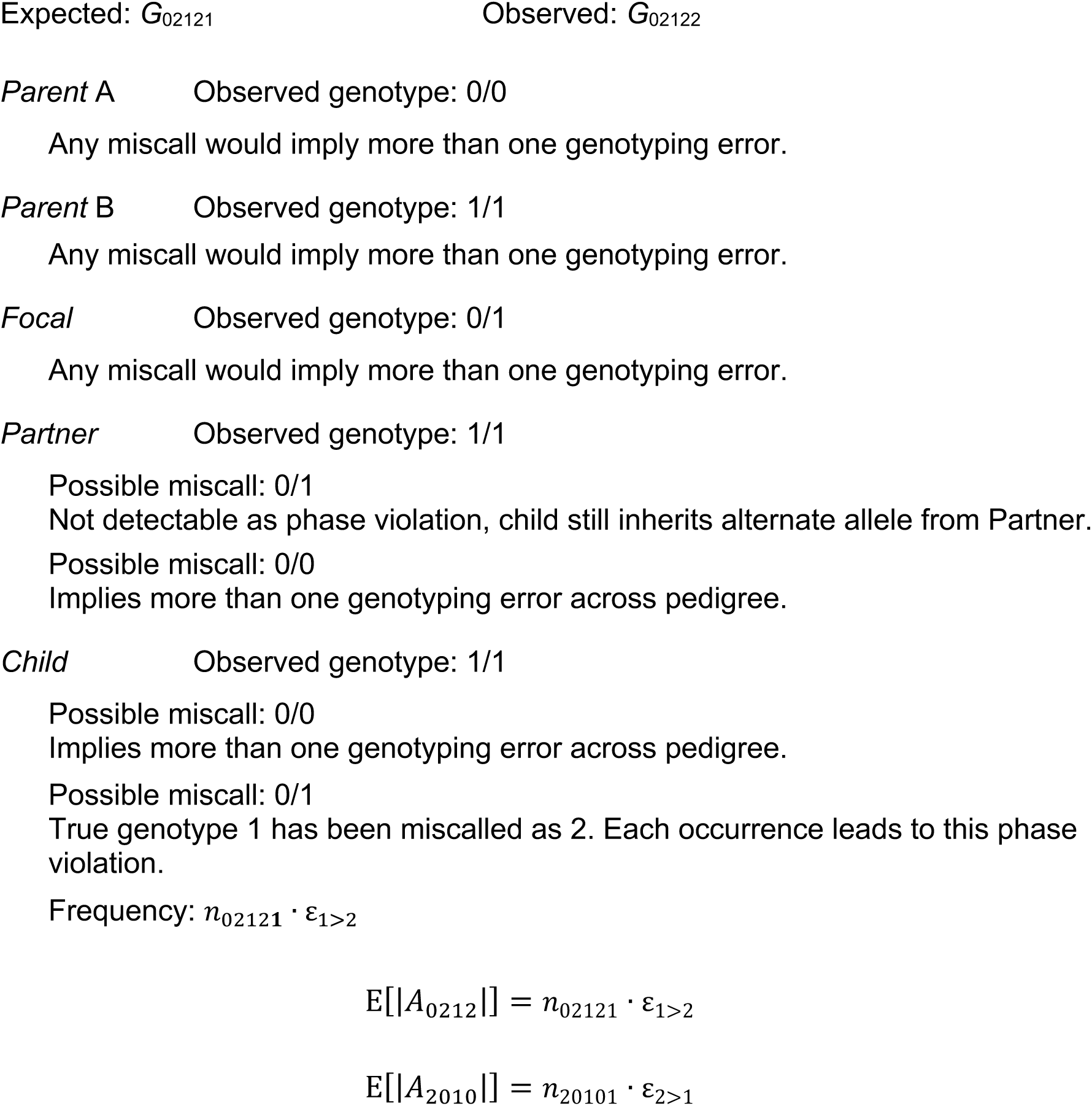

*Expected number of phase violations A*_1212_, *A*_1010_

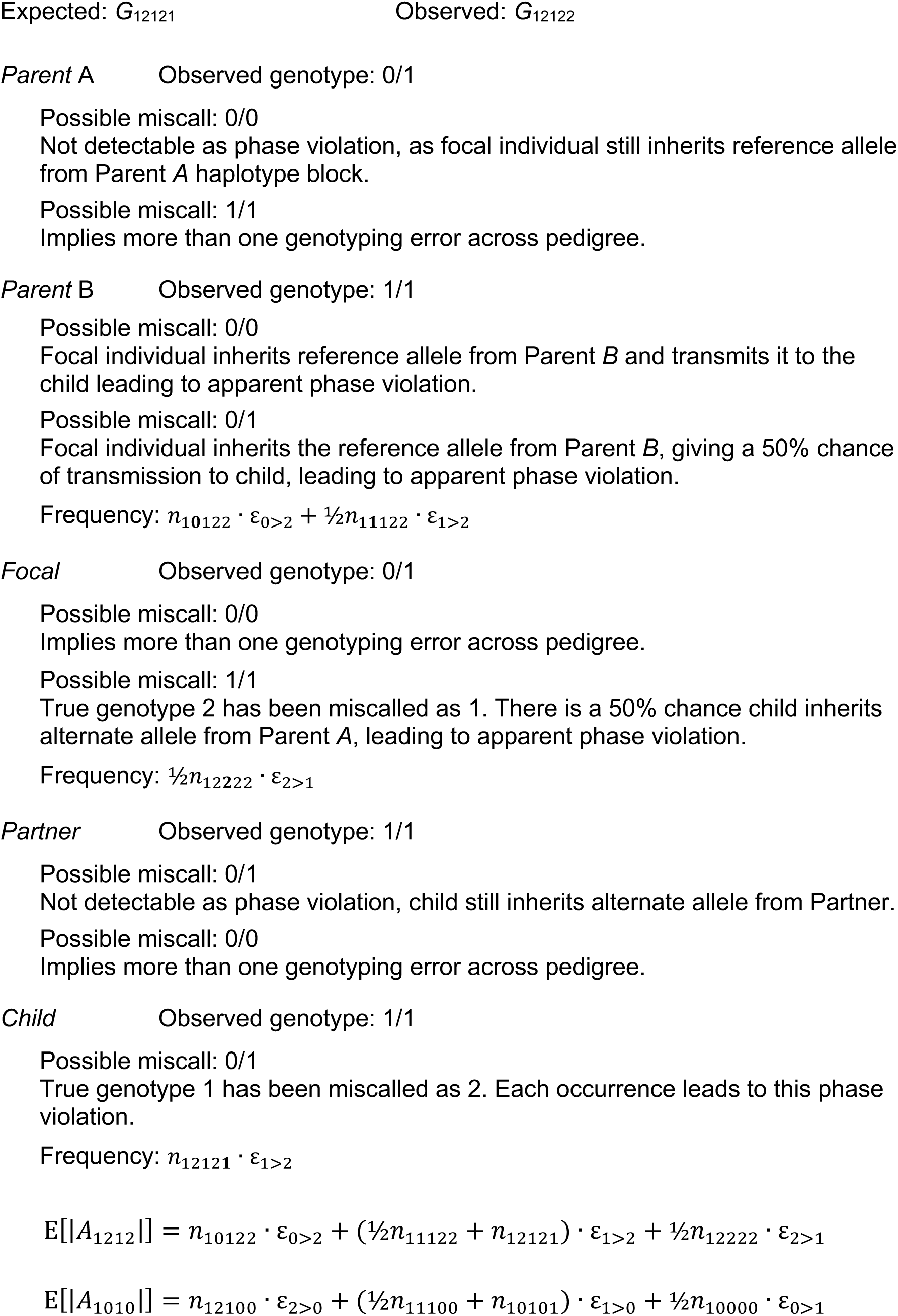

*Expected number of phase violations A*_1211_, *A*_1011_

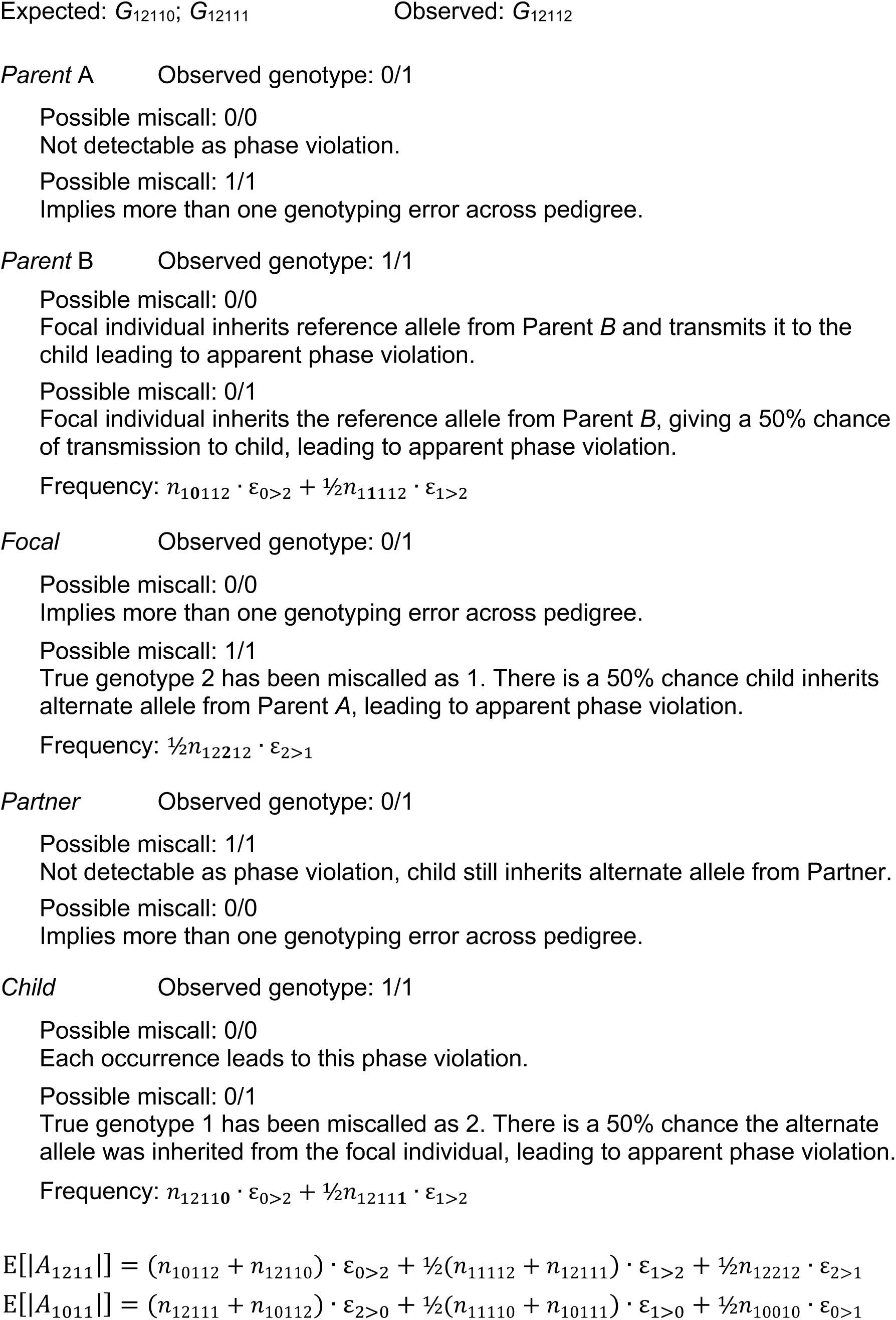

*Expected number of phase violations A*_121*0*_, *A*_101*2*_

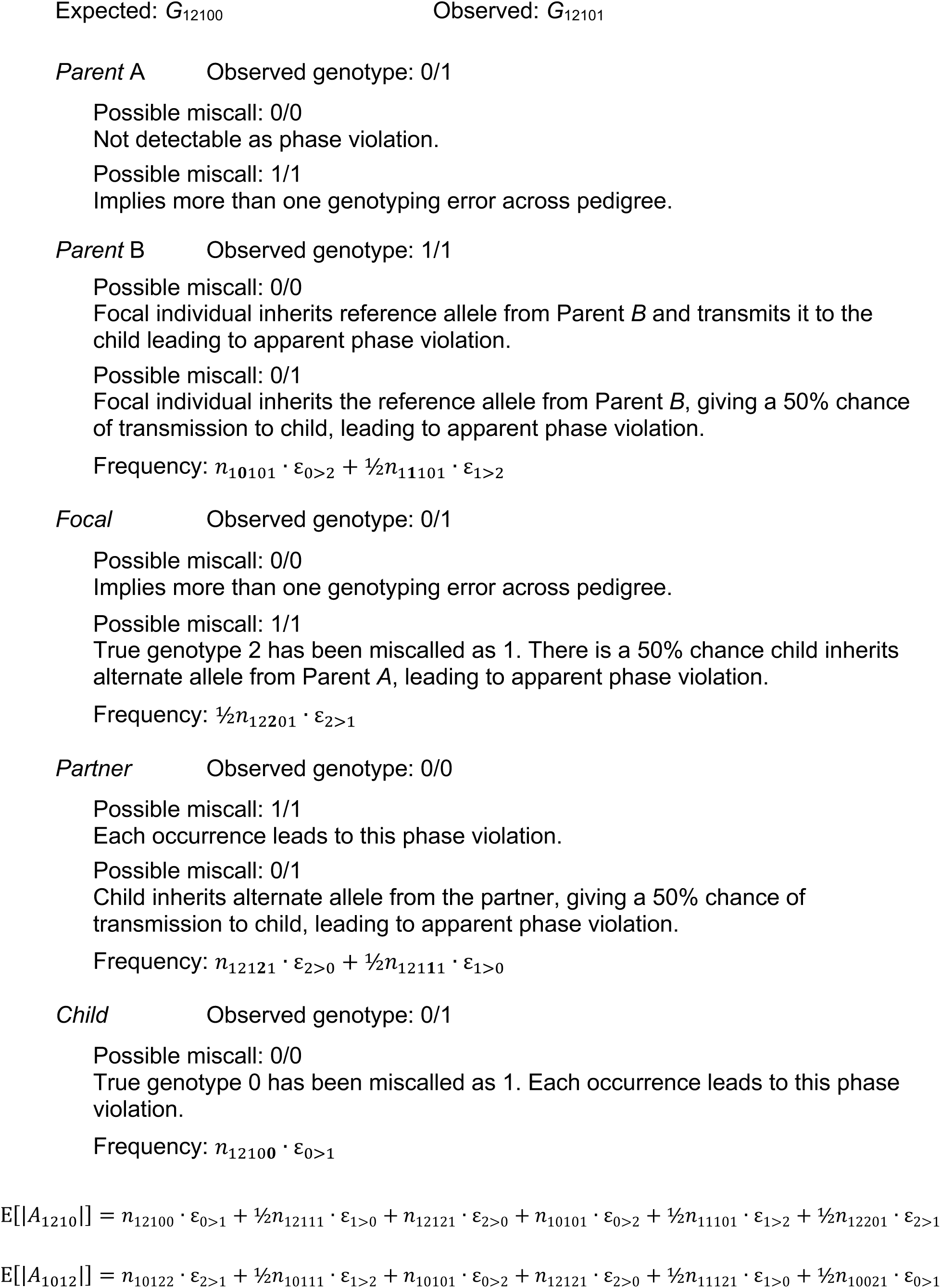

